# Single-cell quantification of the concentration and dissociation constant of endogenous proteins

**DOI:** 10.1101/236422

**Authors:** Akira T. Komatsubara, Yuhei Goto, Yohei Kondo, Michiyuki Matsuda, Kazuhiro Aoki

## Abstract

Kinetic simulation is a useful approach for the elucidation of complex cell-signaling systems. To perform the numerical simulations required for kinetic modeling, parameters such as the protein concentration and dissociation constant (*K_d_*) are essential. However, only a limited number of parameters have been measured experimentally in living cells. Here, we describe a method for quantifying the concentration and *K_d_* of endogenous proteins at the single-cell level with CRISPR/Cas9-mediated knock-in and fluorescence cross-correlation spectroscopy (FCCS). First, the *mEGFP* gene was knocked-in at the end of the *MAPK1* gene, which encoded ERK2, through homology-directed repair or microhomology-mediated end joining. Next, the *HaloTag* gene was further knocked-in at the end of the *RSK2* gene. Protein concentrations of endogenous ERK2-mEGFP and RSK2-HaloTag were quantified in living cells by fluorescence correlation spectroscopy (FCS), revealing substantial cellular heterogeneities. Interestingly, the levels of ERK2-mEGFP and RSK2-HaloTag were strongly positively correlated, suggesting a global mechanism underlying their expressions. In addition, FCCS measurement revealed temporal changes in the apparent *K_d_* values of the binding between ERK2-mEGFP and RSK2-HaloTag in response to EGF stimulation. Our method provides a new approach for quantification of the endogenous protein concentration and dissociation constant in living cells.

---

In response to extracellular signals, mammalian cells process information through an intracellular signaling network comprised of chemical reactions, eventually leading to decisions on cell fate. Extensive studies of cell signaling have identified numerous proteins, pathways, and feedback or feedforward regulations, and have expanded our understanding beyond the simple view of the linear signaling cascade (1). Computer-assisted systems biology approaches may provide a promising strategy for the comprehensive understanding of such complicated cell signaling networks (2). Indeed, various simulation models of signal transduction pathways have been developed in a series of studies over the past 10 years (3–5). Nevertheless, the numerical simulations in most of these systems were conducted with kinetic parameters that have not been measured experimentally, and thus these parameters tend to differ among models, even when the reactions themselves are identical. For this reason, experimentally determined parameters are essential for the development of quantitative and reliable simulation models.

The parameters for simulation of cell signaling are roughly classified into four categories: protein concentration, dissociation constant, diffusion coefficient or transport rate, and enzymatic reaction rate parameters. Classically, these parameters have been measured by *in vitro* biochemical analyses, which require a large number of cells or molecules (6, 7). However, some of these parameters, e.g., protein concentration, are known to show non-genetic cell-to-cell variability (8, 9). Moreover, the quantities of some parameters might differ significantly between *in vitro* and *in vivo*. For instance, we have shown that ERK MAP kinase represents entirely different phosphorylation patterns, namely processive and distributive phosphorylation, under an intracellular and an *in vitro* environment, respectively (7, 10). Therefore, it is of critical importance to measure parameters in living cells.

Fluorescence correlation spectroscopy (FCS) and fluorescence cross-correlation spectroscopy (FCCS) are techniques that exploit the fluctuations of fluorescent molecules in the confocal volume (~1 fL) (11–13). FCS examines the auto-correlation function of temporal fluorescence fluctuation, enabling us to determine the number of fluorescent molecules in the confocal volume and diffusion coefficient. In FCCS, the cross-correlation function is calculated between fluctuations of two different fluorescent species in order to quantify the extent to which these two species form a complex. Thus, FCS and FCCS are capable of measuring *in vivo K_d_* values in living cells, referred to as *in vivo K_d_*. We have previously applied FCS and FCCS to mammalian cells exogenously expressing target proteins used with EGFP and HaloTag for the measurement of *in vivo K_d_* (14). Intriguingly, we found that the *in vivo K_d_* values measured by FCCS were substantially higher than the *in vitro K_d_* values by an order of 1 or 2, because of the competitive binding to non-fluorescently labeled proteins including endogenous and other interacting proteins (14). In addition to the competitive binding, molecular crowding in the cellular milieu has a profound influence on molecular interactions through an excluded-volume effect (15). Thus, the *in vivo K_d_* value reflects the strength of protein-protein interaction within the cells, and would be advantageous in computer simulation of signal transduction, albeit with a variability among different cell types.

It has been reported that the endogenous protein concentration and *in vivo K_d_* value were successfully measured in budding yeast with FCS and FCCS (16, 17). However, there have been no reports of the *in vivo K_d_* values based on the measurement of endogenous proteins in mammalian cells, mainly because of the technical difficulties of knock-in (KI) of a fluorescent protein gene to label the protein of interest.

Recent advances in genome-editing tools have paved the way for tagging endogenous proteins with fluorescent proteins. These genome-editing tools, such as the CRISPR/Cas9 system, enable KI of a gene of interest through DNA double-strand break (DSB) repair mechanisms (18, 19). Homology-directed repair (HDR) is a mechanism by which a homologous template is used as a source of DNA repair. On the other hand, microhomology-mediated end joining (MMEJ) is a mechanism of alternative non-homologous end joining (NHEJ), which also seals DSBs. In contrast to classical NHEJ, MMEJ repairs DNA DSBs using a 5-25 base pair (bp) microhomologous sequence (20). HDR uses a relatively longer homologous sequence (0.1-10 kbp) to seal the DSB, thereby ensuring an error-free repair. Nonetheless, even though MMEJ is an error-prone process of end joining, a recent study demonstrated that MMEJ-mediated KI was more efficient than HDR (21).

In this study, we demonstrate a new approach to quantifying the concentration and *K_d_* of endogenous proteins by combining FCS and FCCS with CRISPR/Cas9-mediated KI of fluorescent proteins in mammalian cells.

## Results

### Design of donor vectors and KI strategy

First, we attempted to knock-in the *mEGFP* gene at the 3’ site of the human *MAPK1* gene encoding ERK2 in HeLa cells. To do this, we constructed a selection cassette for the donor vector, which contained the *mEGFP* gene, a loxP sequence, a porcine teschovirus-1-derived self-cleaving 2A peptide (P2A) sequence, a bi-functional fusion protein between a truncated version of herpes simplex virus type 1 thymidine kinase (*dTK*) and the bacterial neomycin phosphotransferase (*neo*) genes (*dTKneo*), a poly(A) addition sequence, and a loxP sequence (Fig. 1A and Fig. S1). To develop a donor vector for KI at the *hMAPK1* gene, this cassette was further sandwiched between the longer homology arms (L. arm and R. arm) or the shorter homology arms (40 bp for each) for HDR- or MMEJ-mediated DSB repair (Fig. 1A, top and bottom). The *dTKneo* was generated based on previous reports (22, 23) as a positive/negative selection marker, and connected to *mEGFP* via cDNA of the P2A sequence, which is a self-cleaving peptide so that genes sandwiching the P2A peptide are separately expressed (24). Therefore, the expression of dTKneo requires both in-frame integration and endogenous promoter activity of the gene prior to the KI cassette, resulting in the suppression of false-positive cells harboring random insertions of the cassette. dTKneo expression renders cells resistant to G418 and sensitive to Ganciclovir. Of note, it has been reported that *neo* is preferable for targeting moderate or low expression genes with the promoter-less targeting vector (25). After KI of the donor vector, Cre-mediated recombination removes the P2A, dTKneo and poly(A) addition sequence. Ganciclovir treatment selects the cells whose selection cassette has been removed successfully (Fig. 1A).

**Figure 1.**
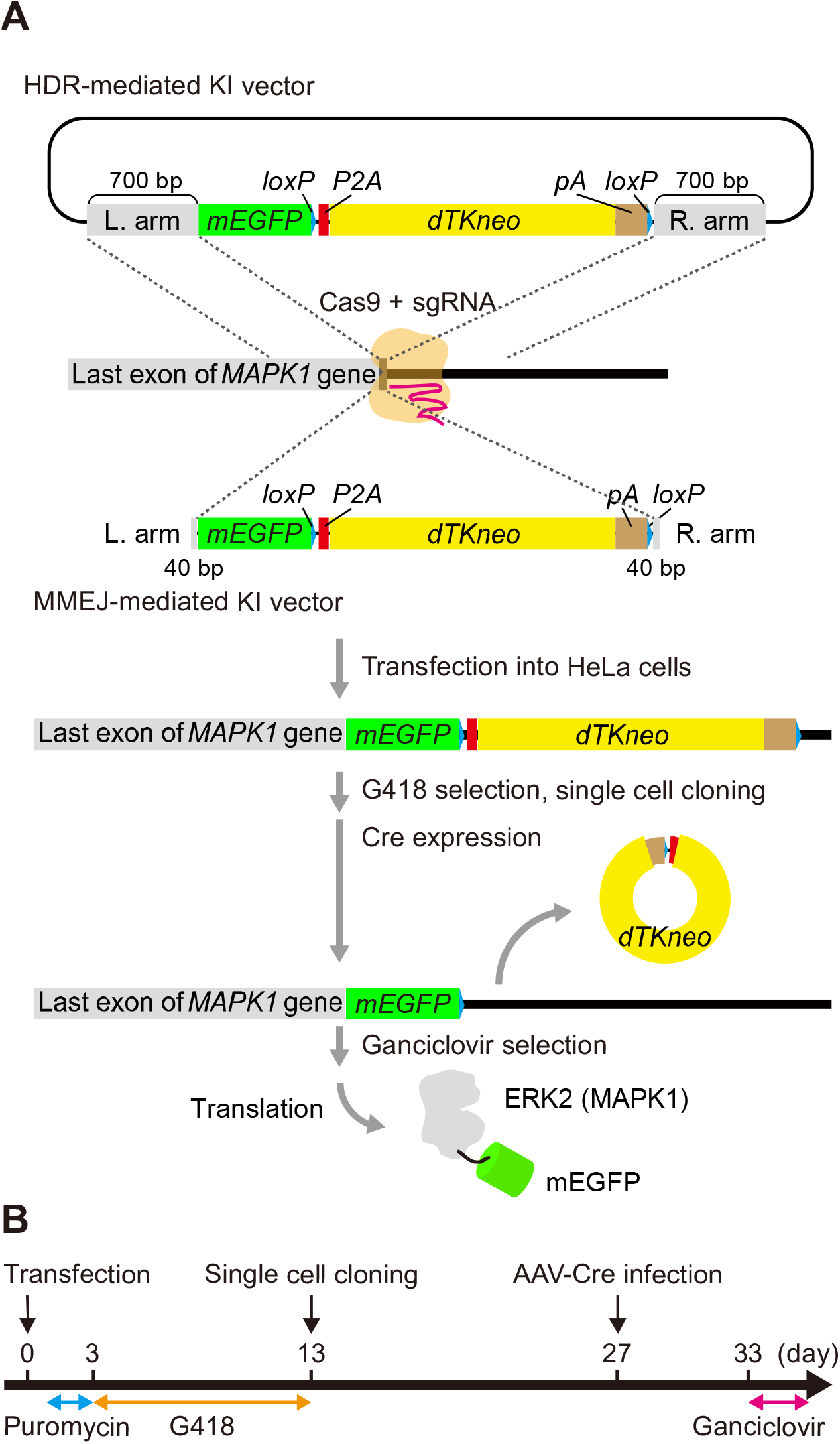
Outlines of gene KI. (A) Schematic illustration of gene KI with an HDR-mediated or MMEJ-mediated KI vector at the human *MAPK1* locus. (B) Procedure for *mEGFP* KI. HeLa cells were transfected with the donor vector and Cas9 vector at day 0, and transfected cells were selected by puromycin. The cells were then subjected to G-418 selection for approximately 10 days. After G-418 selection, each GFP-positive cell was sorted into a 96-well plate by a FACSAria IIu flow cytometer (BD Biosciences). Single clonal cells were infected with AAV-Cre to remove the P2A-dTKneo-polyA (pA) cassette.

In our experimental schedule, we transfected the pX459 CRISPR/Cas9 vector and donor DNA into HeLa cells, then selected the transfected cells by 1.0 μg/mL puromycin treatment. Three days after transfection, we began to select KI cells with 0.5 mg/ml G418. Thirteen days after selection, the KI cells were sorted by flow cytometry based on mEGFP fluorescence, followed by single cell cloning for an additional 14 days. The cells were then infected with adeno-associated virus (AAV), which transiently induced the expression of Cre recombinase, to remove the P2A-dTKneo cassette. One week after AAV infection, the cells were further subjected to negative selection with Ganciclovir.

### Establishment of KI HeLa cells expressing ERK2-mEGFP

As a proof-of-concept, we targeted the *MAPK1* gene to generate cells expressing ERK2-mEGFP from the endogenous locus. HeLa cells were transfected with pX459-hMAPK1 and a donor vector for HDR- or MMEJ-mediated KI (Fig. 1A). After G418 selection, the genomic DNA were extracted from parental HeLa cells or bulk HeLa cells transfected with pX459-hMAPK1 and HDR- or MMEJ-donor vector, and subjected to PCR with either the forward (F) primer, the reverse (R) primer or both primers (FR). As expected, PCR products were only observed in the sample from cells introduced with the HDR donor or MMEJ donor when we used both the forward and reverse (FR) primers (Fig. 2A), indicating KI of donor vectors into the 3’ site of the *MAPK1* gene.

**Figure 2.**
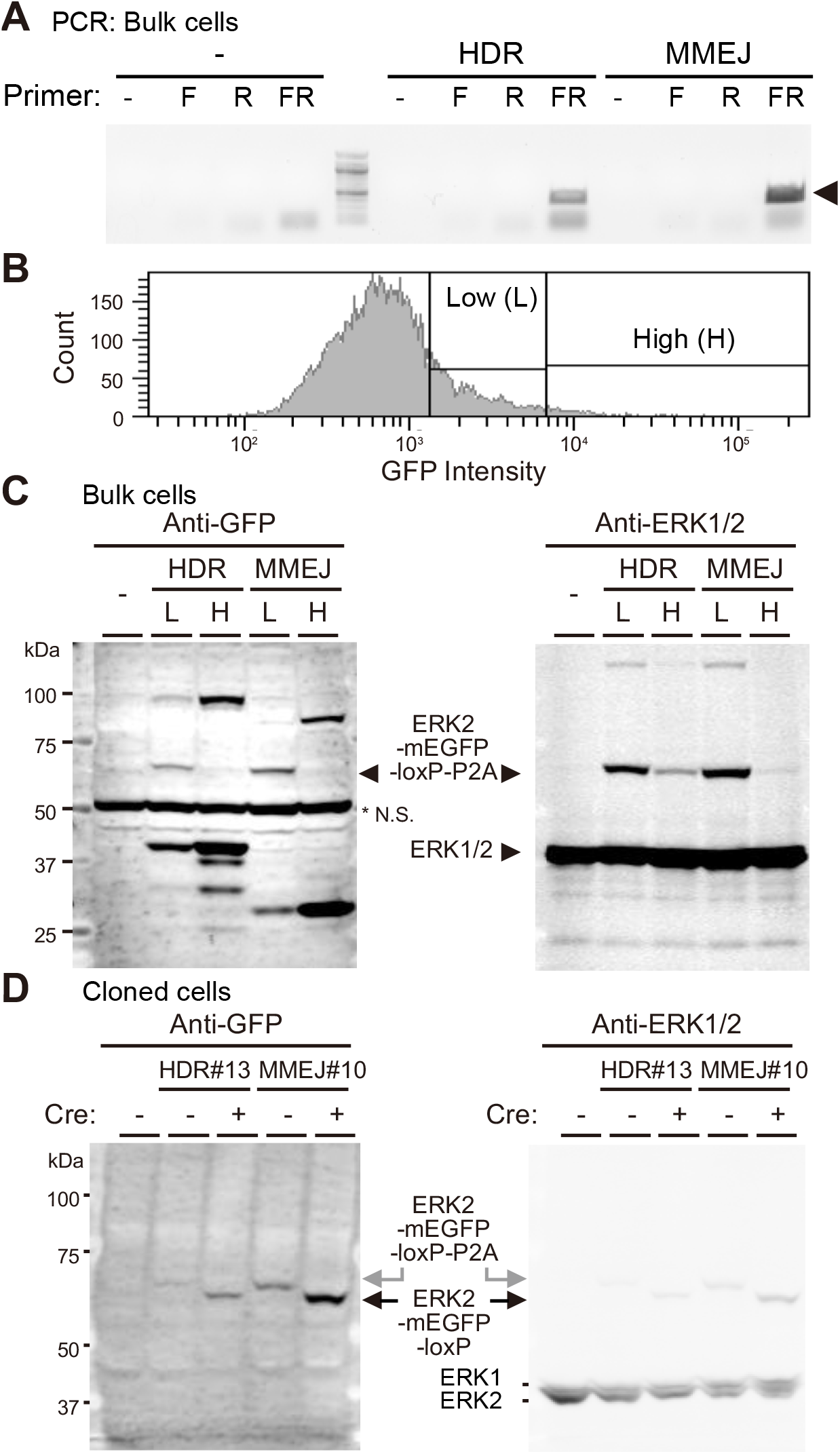
*mEGFP* KI at the human *MAPK1* locus. (A) After G-418 selection, the genomic DNAs of bulk HeLa cells were extracted and analyzed by PCR to confirm *mEGFP* integration. -, F and R represent the negative control, forward primer and reverse primer, respectively. The forward primer and the reverse primer bind to *MAPK1* and *mEGFP*, respectively. The arrowhead indicates the expected PCR products. (B) Histograms showing the distribution of GFP intensity of bulk HeLa cells, in which the *mEGFP* gene was integrated via MMEJ. GFP-positive cells were divided into Low (L) and High (H) groups. (C) The cell lysates were obtained from parental HeLa cells (−), Low (L) and High (H) cells in panel B, and subjected to immunoblotting with anti-GFP (left) and anti-ERK1/2 (right) antibodies. The arrowhead indicates ERK2-mEGFP. An asterisk (*) indicates a non-specific (N.S.) signal. (D) *mEGFP* knocked-in HeLa cells by HDR-mediated or MMEJ-mediated KI vector were subjected to single cell cloning. The cell lysates from HeLa/ERK2-mEGFP-HDR#13 and HeLa/ERK2-mEGFP-MMEJ#10 cells before (-) and after (+) AAV-Cre infection were analyzed by immunoblotting with anti-GFP (left) and anti-ERK1/2 (right) antibodies.

Next, we used flow cytometry to sort the transfected HeLa cells into two populations based on their GFP fluorescence intensities: a low GFP fluorescence (Low) and high GFP fluorescence (High) group (Fig. 2B). The Low and High cells accounted for 17.2% and 2.0% of the parent population in HeLa cells transfected with HDR-donor vector, respectively (Fig. 2B). Meanwhile, 5.6% and 0.1% of the parent population in cells transfected with the MMEJ-donor vector were categorized as Low and High cells, respectively. The results of immunoblotting with an anti-GFP antibody and an anti-ERK1/2 antibody demonstrated the expression of ERK2-mEGFP, with a molecular weight of approximately 69 kDa (arrowhead), in both Low and High cells transfected with HDR- or MMEJ-vector (Fig. 2C). Intriguingly, in both HDR- and MMEJ-mediated KI, High cells showed stronger non-specific mEGFP-integration bands and weaker specific ERK2-mEGFP-integration bands than Low cells (Fig. 2C).

We subsequently isolated several single clones of Low cells by the limiting dilution method. Immunoblotting with the anti-ERK1/2 antibody demonstrated that isolated cell lines expressed comparable levels of ERK2-mEGFP (Fig. S2A). Western blot analysis revealed that, in comparison to parental HeLa cells, some of the isolated cells showed only a positive signal of ERK2-mEGFP around 69 kDa (cell lines #13, #16, and #20 with the HDR-donor vector and cell lines #1, #3, #5, #10, and #12 with the MMEJ-donor vector). The other cells exhibited not only the positive ERK2-mEGFP signal but also apparently negative GFP signals, indicating higher or lower molecular weights than we expected (Fig. S2A). Thus, we picked up the former cell clones, and confirmed that the positive ERK2-mEGFP signals were also detected with anti-ERK2 antibody (Fig. S2B). The KI efficiencies were roughly 15% (3 out of 20) and 40% (5 out of 12) with the HDR- and MMEJ-donor vectors, respectively (Fig. S2).

For further analysis, HDR-mediated KI cell line #13 (HDR #13) and MMEJ-mediated KI cell line #10 (MMEJ #10) were infected with AAV expressing Cre recombinase, followed by the analysis of expression levels of ERK2-mEGFP (Fig. 2D). Removal of the P2A-dTKneo-poly(A) cassette increased the ERK2-mEGFP levels in both cell lines (2.54-fold increase) (Fig. 2D, left). The expression level of endogenous ERK2 in control cells was equal to an 8.05-fold increase in the expression of ERK2-mEGFP (MMEJ #10). This might be due to the transfer efficiency during the experimental process and/or the effect of fusion of mEGFP to ERK2 at the C-terminus. Of note, the expression levels of endogenous ERK2 did not show a discrete reduction, i.e., a 50% or 100% reduction, but rather exhibited a 40% to 80% reduction in the KI cell lines (Fig. 2D, right), suggesting that *mEGFP* genes were knocked-in into one or two of the multiple *MAPK1* gene loci in HeLa cells, which are known to demonstrate aneuploidy (26).

### Establishment of RSK2-HaloTag KI HeLa cells

Because RSK2 is known to interact with ERK2 (27), we next targeted the *RSK2* gene for labeling with HaloTag, which covalently linked to HaloTag ligands such as the tetramethylrhodamine (TMR)-ligand (28). For *HaloTag* gene integration, we tested the aforementioned MMEJ-donor and precise integration into the target chromosome (PITCh)-like KI vector (21)(Fig. 3A). The PITCh-vector includes the insertion cassette, which is further sandwiched by recognition sites of the Cas9/gRNA complex that target the *RSK2* gene locus (Fig. 3A, lower). Therefore, the Cas9/gRNA complex digests not only the RSK2 gene locus but also the PITCh-like donor vector, generating linear double strand donors within the cells.

**Figure 3.**
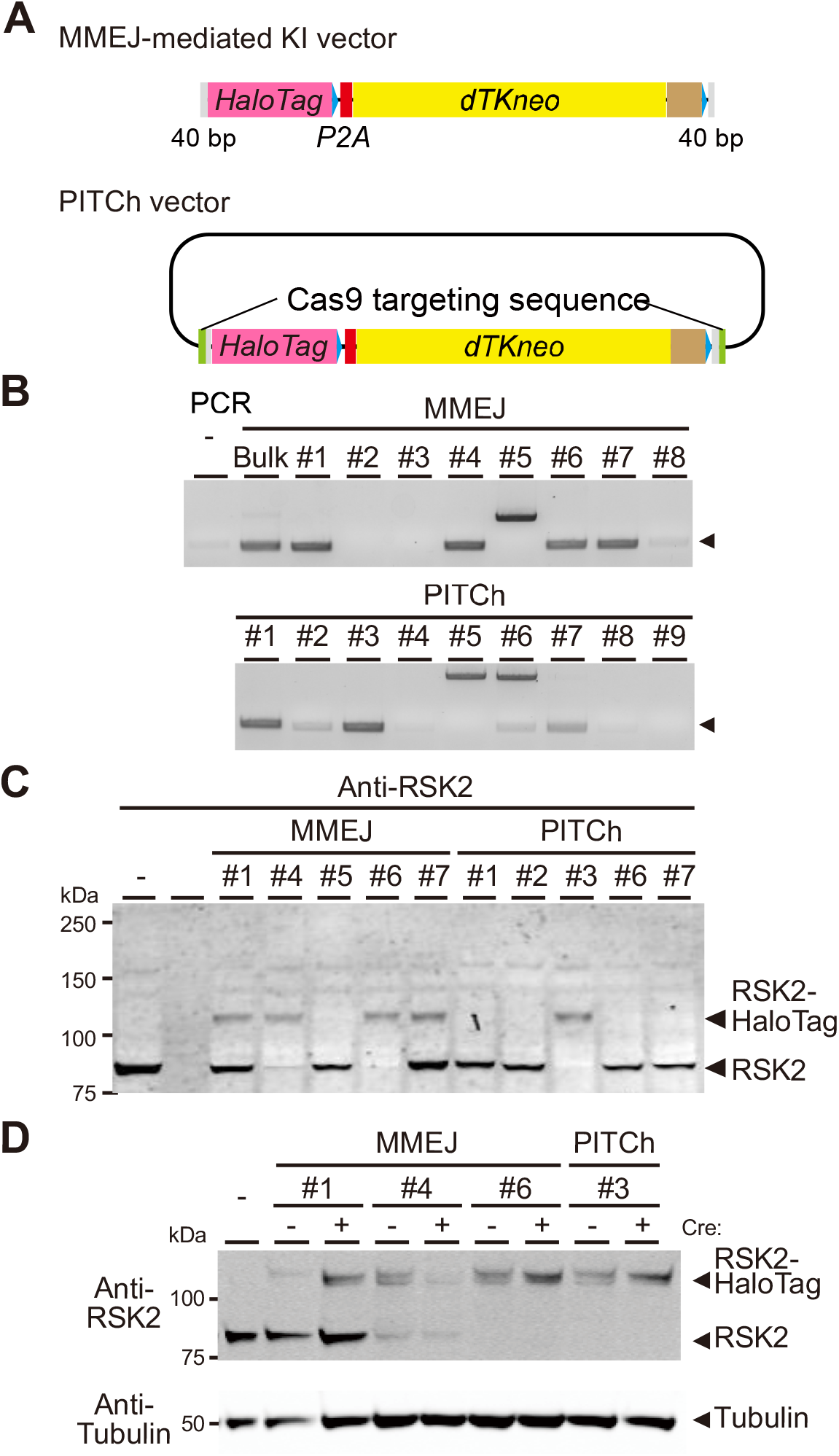
*HaloTag* KI at the human *RSK2* locus. (A) Schematic representation of the MMEJ-mediated KI vector (upper) and PITCh-like KI vector (lower) for *HaloTag* gene KI. The PITCh vector contains the sequence targeted by sgRNA, which recognizes the *RSK2* gene. (B) After G-418 selection, genomic DNAs were extracted, and analyzed by PCR to confirm *HaloTag* integration. Eight and nine clones were analyzed in clonal HeLa cells transfected with MMEJ-mediated KI vector (MMEJ, upper) and PITCh vector (PITCh, lower), respectively. The forward primer and the reverse primer bind to *RSK2* and *mHaloTag*, respectively. The arrowhead indicates the expected PCR products. (C) The cell lysates were obtained from parental HeLa cells (-), and the indicated clones of HeLa/RSK2-HaloTag-MMEJ (MMEJ) and HeLa/RSK2-HaloTag-PITCh (PITCh) cells in panel B, and subjected to immunoblotting with anti-RSK2 antibody. (D) The cell lysates were obtained from parental HeLa cells (-), and the indicated clones of HeLa/RSK2-HaloTag-MMEJ (MMEJ) and HeLa/RSK2-HaloTag-PITCh (PITCh) cells before (-) or after (+) AAV-Cre infection, and subjected to immunoblotting with anti-RSK2 (upper) and anti-tubulin (lower) antibodies.

To obtain HeLa cells expressing ERK2- mEGFP and RSK2-HaloTag from this endogenous locus, we introduced Cas9 and the sgRNA expression vector (pX459-hRSK2), which targeted a site close to the stop codon of the *RSK2* gene, and linearized the MMEJ-donor vector (Fig. 3A, upper) or the PITCh-like donor plasmid (Fig. 3A, lower) into HeLa/ERK2- mEGFP (MMEJ#10) cells (Fig. 2D). After G418 selection and single cell cloning, several clones were analyzed by PCR using two primers that recognized the 3’ end of *RSK2* or the 5’ end of *HaloTag* (Fig. 3B). Almost half of the clones used in either donor vector showed the expected PCR product (Fig. 3B). However, subsequent DNA sequencing of these PCR products revealed the repeated sequence of the homology arm in PITCh #1, 2, 6, and 7 cells, resulting in a frame shift. Meanwhile, MMEJ #1, 4, 6, and 7 cells had no insertion or deletion. In agreement with these data, the expressions of RSK2-HaloTag were observed only in MMEJ #1, 4, 6, 7 and PTICh #3 cells by immunoblotting with anti-RSK2 antibody (Fig. 3C). As with the case of ERK2-mEGFP, the removal of the selection marker also enhanced the expression of the RSK2-HaloTag (1.89-fold increase) except in MMEJ #4 cells (Fig. 3D). The expression level of the RSK2-HaloTag was 5.14fold lower than the expression of RSK2 in the control cells (Fig. 3D).

### Subcellular localization of ERK2-mEGFP and RSK2-HaloTag KI HeLa cells

It is well-known that mitogen stimulation induces nuclear accumulation of ERK, thereby causing phosphorylation of several transcription factors and subsequent gene expression (29–31). We picked up the cell line PITCh #3 (HeLa/ERK2-mEGFP/RSK2-HaloTag), and visualized the RSK2-HaloTag by staining with 100 nM TMR-Ligand. As expected, treatment with fetal bovine serum (FBS) and epidermal growth factor (EGF) provoked nuclear translocation of ERK2-EGFP from the cytoplasm, but did not induce subcellular translocation of RSK2-HaloTag-TMR within 30 min (Fig. 4A and 4B). The kinetics of nuclear translocation of ERK2-mEGFP was slightly delayed compared with that of endogenous ERK1/2 observed by immunofluorescence (Fig. S3A and S3B), suggesting that fusion with mEGFP had a negative effect on the nucleocytoplasmic shuttling of ERK2. RSK2-HaloTag-TMR showed nuclear translocation by FBS stimulation for 4 h (Fig. S3C and S3D), consistent with the previous study (32). Fluorescence speckles in KI cells were also observed in parental HeLa cells under our experimental condition (Fig. S3E). Taken together, these data provide evidence that ERK2-mEGFP and RSK2-HaloTag can be imaged by fluorescence microscopy.

**Figure 4.**
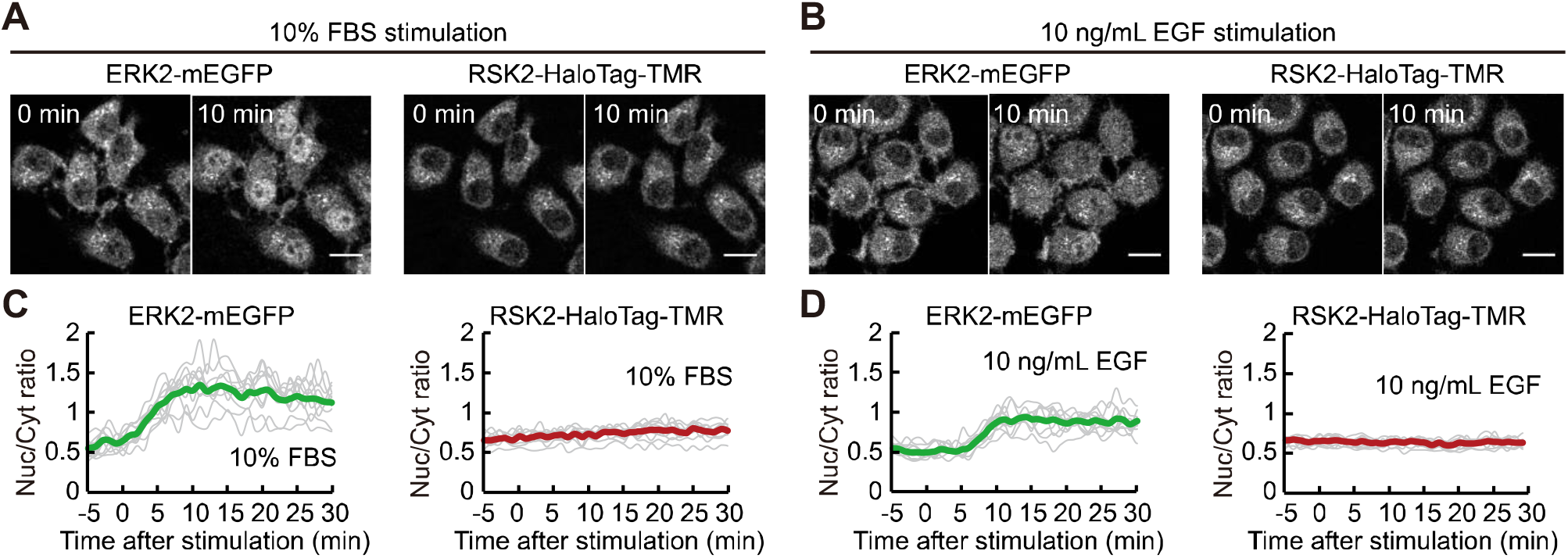
Subcellular distribution of ERK2 and RSK2 in HeLa cells stimulated with EGF or FBS. HeLa/ERK2-mEGFP-MMEJ#10/HaloTag-PITCh#3 cells were stained with tetramethylrhodamine (TMR)-ligand and stimulated with 10% FBS (A and C) or 10 ng/mL EGF (B and D). (A and B) Representative images of ERK2-mEGFP (left) and RSK2-HaloTag-TMR (right) are shown before and 10 min after stimulation. Scale bars, 20 nm. (C and D) Time course of the ratio of nuclear to cytosol fluorescence in ERK2-mEGFP (left) and RSK2-HaloTag-TMR (right) are plotted (N = 10 cells for all conditions). Thin lines and thick lines indicate the time course data obtained from each cell and the average value, respectively.

### Evaluation of measurement noise and intra- and intercellular variability of protein concentration measured by FCS

To quantitatively address the efficacy of endogenous protein measurement by FCS, we assessed the variability derived from labeling efficiency, measurement noise, and intra- and intercellular heterogeneity. First, we confirmed that the HaloTag-TMR-ligand was completely labeled throughout the more than 6 h of incubation under our experimental condition (Fig. 5A). HaloTag protein is known to covalently bind to HaloTag-ligand, and thus we concluded that HaloTag proteins exhibited almost 100% labeling efficiency. Under this condition, mEGFP-HaloTag-TMR fusion proteins were transiently expressed in HeLa cells, and analyzed by FCS (Fig. 5B). The number of mEGFP proteins obtained by FCS was plotted as a function of the number of HaloTag-TMR proteins, showing a linear correlation with a slope of 0.92 (Fig. 5C). If we assume 100% labeling efficiency of HaloTag-TMR, we can derive 92% of the maturation efficiency of the mEGFP fluorophore.

**Figure 5.**
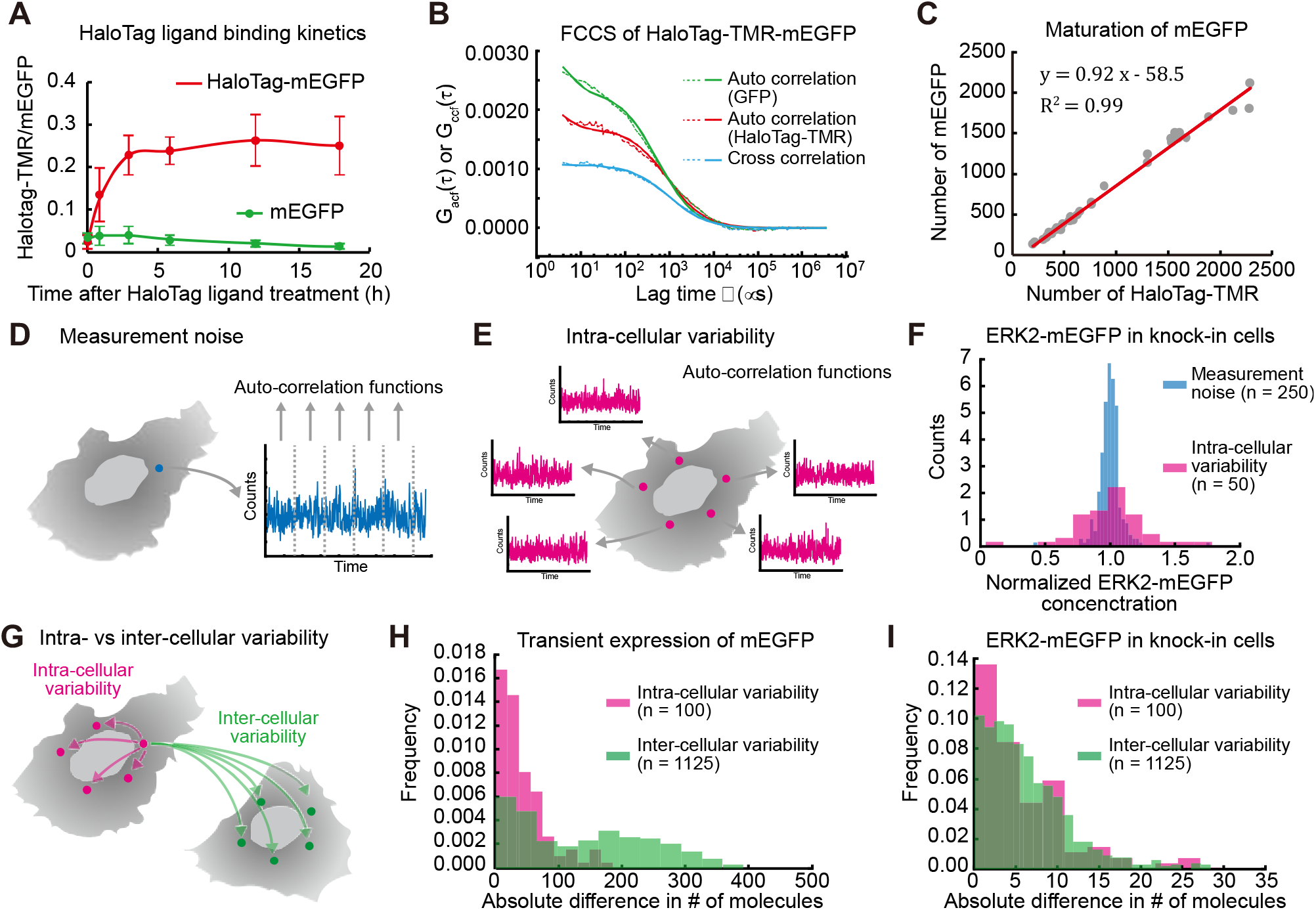
Evaluation of measurement noise, and intra- and inter-cellular variability in FCS analysis. (A) HeLa cells expressing mEGFP (green) or HaloTag-mEGFP (red) were labeled with TMR-ligand for the indicated amount of time. Averaged HaloTag-TMR/mEGFP ratio values are plotted with SD (N > 50 cells for all conditions). (B and C) HeLa cells expressing HaloTag-TMR-mEGFP were analyzed by FCS and FCCS. Representative correlation curves are shown. Thin and bold lines show the experimental data and fitting curves, respectively. (B). The numbers of mEGFP molecules are plotted as a function of the number of HaloTag-TMR molecules with a fitted line (red) (C). (D and E) Schematic representation of the evaluation of measurement noise (D) and intra-cellular variability (E) in FCS analysis. (F) ERK2-mEGFP concentration in HeLa/ERK2-mEGFP-MMEJ#10/HaloTag-PITCh#3 was measured by FCS, and the normalized ERK2-mEGFP concentration was calculated as in D and E. The histograms of normalized ERK2-mEGFP obtained by C and D are shown. (G) Schematic representation of the comparison of intra-cellular variability with inter-cellular variability. (H) HeLa cells transiently expressing mEGFP were analyzed by FCS. The differences in the number of mEGFP molecules were calculated as in G, and plotted as a histogram. (I) ERK2-mEGFP concentration in HeLa/ERK2-mEGFP-MMEJ#10/HaloTag-PITCh#3 was measured by FCS. The differences in the number of mEGFP molecules were calculated as in G, and plotted as a histogram.

Next, we compared the measurement noise of FCS with intracellular variability. To calculate measurement noise, single-point FCS time series data were divided into 5 segments, and each segment was separately analyzed to obtain auto-correlation functions and protein concentrations (Fig. 5D). For intracellular variability, we picked up 5 different points for FCS measurement in a single cell, and obtained protein concentrations at these 5 points (Fig. 5E). The results showed that the measurement noise was much smaller than the intracellular variability (Fig. 5F).

To further evaluate the difference between the intracellular and intercellular variability, we calculated the absolute difference of the number of molecules between two points (Fig. 5G). As a positive control, we transiently overexpressed mEGFP-HaloTag proteins in HeLa cells, measured the number of mEGFP molecules by FCS, and calculated the differences of intra- and intercellular variability. As expected, the histogram of the inter-cellular variability showed a broader distribution of the absolute difference of the number of molecules than the intra-cellular variability (Fig. 5H), indicating that this method is able to discriminate between intra- and intercellular variability. Next, we examined the inter-cellular variability in ERK2-mEGFP KI cells. To our surprise, we did not observe any substantial difference in ERK2-mEGFP expression between the intra- and inter-cellular variability (Fig. 5I). This may be because the inter-cellular variability is smaller than the intracellular variability and therefore masked by the intracellular variability. This result was also supported by comparing these variabilities with those measured using a conventional confocal microscope; the intra-cellular variability was larger than the inter-cellular variability in both ERK2-mEGFP and RSK2-HaloTag-TMR (Fig. S4). These results indicated that FCS was not well suited to assessing the inter-cellular variability in ERK2-mEGFP and RSK2-HaloTag-TMR.

### Quantification of the endogenous protein concentration and in vivo K_d_ value by FCS and FCCS

Finally, we measured the protein concentration and the *in vivo K_d_* values of ERK2-mEGFP and RSK2-HaloTag in HeLa/ERK2-mEGFP/RSK2-HaloTag cells by FCS and FCCS, respectively. The auto-correlation functions of ERK2-mEGFP and RSK2-HaloTag-TMR and cross-correlation functions between ERK2-mEGFP and RSK2-HaloTag-TMR were calculated in a single HeLa cell (Fig. 6A). We performed repeated measurements with FCS and FCCS, and obtained the distribution of the endogenous protein concentration of ERK2-mEGFP and RSK2-HaloTag-TMR from 198 cells (Fig. 6B). The average concentrations of ERK2-mEGFP and RSK2-HaloTag-TMR were 0.078 μM and 0.097 μM, respectively (Fig. 6B). Total ERK2 and RSK2 concentrations in a HeLa cell were estimated as 0.68 μM and 0.50 μM, respectively, based on western blotting data (Fig. 2D for ERK2 and Fig. 3D for RSK2) and the fluorophore maturation efficiency of mEGFP (Fig. 5B) (see Experimental Procedures). As mentioned above, the heterogeneity of ERK2 and RSK2 concentration was mainly attributed to the intra-cellular variability (Fig. 5 and Fig. S4). Nevertheless, a positive correlation between ERK2-mEGFP and RSK2-HaloTag-TMR was found, with a correlation coefficient of 0.64 (Fig. 6B).

**Figure 6.**
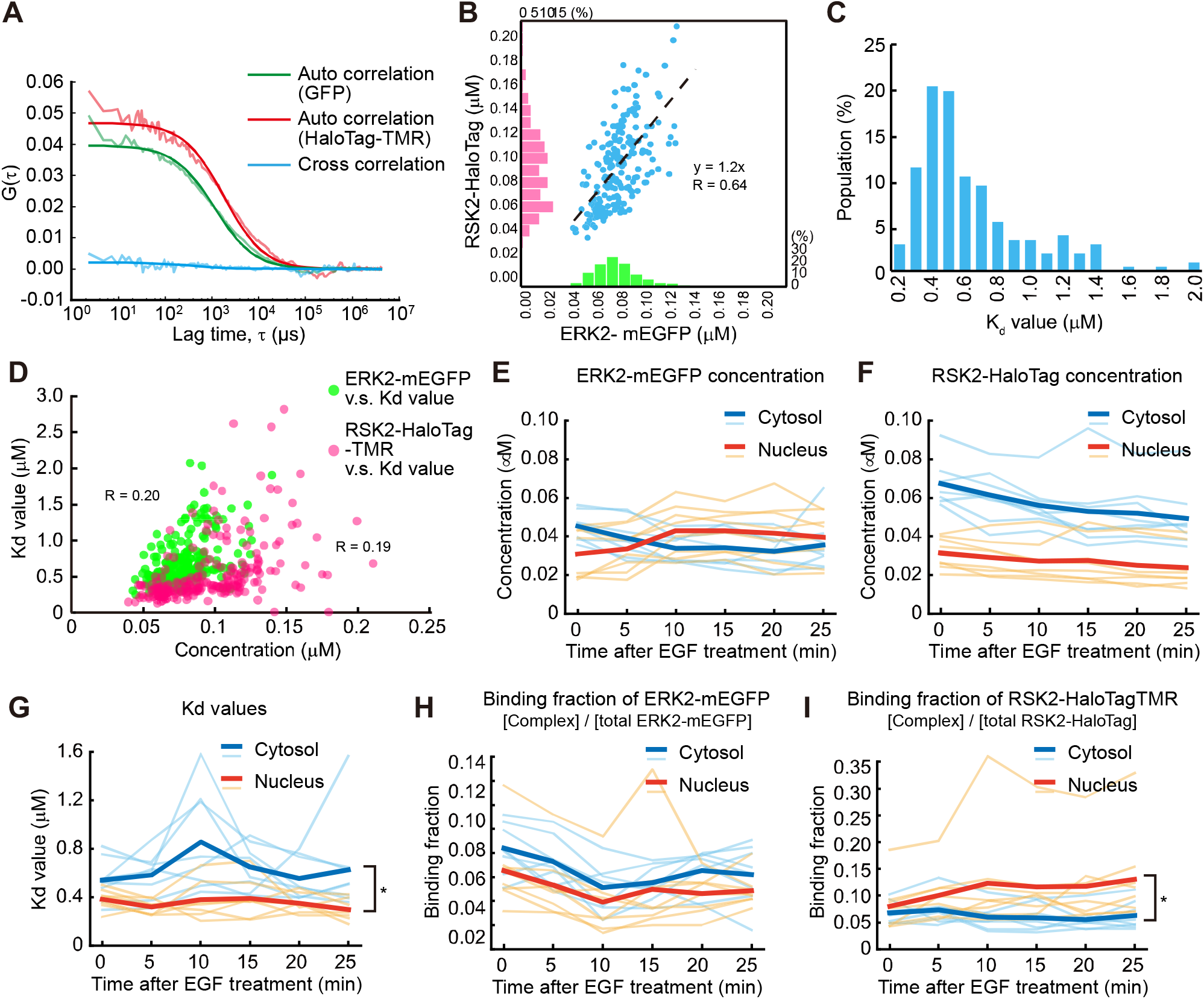
Quantification of endogenous ERK2-mEGFP and RSK2-HaloTag-TMR concentration and *K_d_* values by FCS and FCCS. (A) Representative results of FCS and FCCS in HeLa/ERK2-mEGFP-MMEJ#10/HaloTag-PITCh#3 cells. Thin and bold lines are experimental data and fitting curves, respectively. (B) Scatter plot and histograms of ERK2-mEGFP and RSK2-HaloTag-TMR concentrations in individual HeLa/ERK2-mEGFP-MMEJ#10/HaloTag-PITCh#3 cells (n = 198 from 3 independent experiments). R indicates a correlation coefficient. (C) Histogram of *K_d_* values obtained by FCCS in individual HeLa/ERK2-mEGFP-MMEJ#10/HaloTag-PITCh#3 cells. (D) Scatter plots of ERK2-mEGFP vs *K_d_* values (green dots) and RSK2-HaloTag-TMR vs *K_d_* values (magenta dots). R indicates a correlation coefficient. (E-I) ERK2-mEGFP (E), RSK2-HaloTag-TMR (F) concentration, *K_d_* values (G), binding ratio values of ERK2-mEGFP (H) and RSK2-HaloTag-TMR (I) in the cytosol (blue) and nucleus (red) are plotted as a function of time after EGF treatment. Thin and bold lines indicate individual and averaged data, respectively. N = 8 measurements from 4 independent experiments. \**p <* 0.05 (cytosol vs nucleus) by ANOVA.

The average *in vivo K_d_* value between ERK2-mEGFP and RSK2-HaloTag-TMR was approximately 0.63 μM (Fig. 6C). The *in vivo K_d_* values also showed little correlation with the protein concentrations of ERK2-mEGFP and RSK2-HaloTag-TMR (Fig. 6D). We confirmed that ERK2-mEGFP and RSK2-HaloTag-TMR did not bind nonspecifically to HaloTag-TMR and mEGFP, respectively (Fig. S5).

Finally, in order to take full advantage of this strategy, we quantified the temporal changes in *in vivo K_d_* values at the subcellular level. HeLa/ERK2-mEGFP/RSK2-HaloTag cells were stained with the TMR-HaloTag ligand, then treated with epidermal growth factor (EGF), followed by measurement of the protein concentration and *in vivo K_d_* values in the cytoplasm and nucleus. As we demonstrated in Figure 4, ERK2-mEGFP, but not RSK2-HaloTag-TMR, was translocated from cytoplasm to nucleus upon EGF stimulation (Fig. 6E and 6F). The nuclear concentration of ERK2-mEGFP after EGF stimulation surpassed the cytoplasmic one (Fig. 6E). The *in vivo K_d_* values in the cytosol were higher than those in the nucleus over the course of EGF stimulation (Fig. 6G). In addition, the *K_d_* value in the cytosol, but not that in the nucleus, was transiently increased 10 min after EGF stimulation (Fig. 6G). These results indicated that the ERK2-RSK2 complex was more stable in the nucleus than in the cytosol under both the unstimulated and stimulated conditions, and EGF stimulation transiently reduced the apparent affinity of the ERK2-RSK2 complex only in the cytoplasm. The values of the ratio of complex to ERK2-mEGFP did not show any significant change, because ERK2-mEGFP *per se* shuttled between the cytoplasm and nucleus (Fig. 6H). In contrast, the ratio of complex to RSK2-Halotag-TMR gradually increased in the nucleus (Fig. 6I). This is because the RSK2 concentration did not change and ERK2 flowed into the nucleus.

## Discussion

Here, we established a method for quantifying the endogenous protein concentration and dissociation constant in living cells with a CRISPR/Cas9 genome editing technique and FCS/FCCS. In principle, these techniques allow us to quantify these kinetic parameters without any antibodies. While several research groups have reported methods for endogenous tagging in cultured mammalian cells (21, 33, 34), gene KI is still challenging in cultured cells even when using the CRISPR/Cas9 technique. The KI efficiency is much lower than the random integration efficiency in HeLa cells. Therefore, it is important not only to increase the KI efficiency (35-37), but also to decrease the emergence of negative clones. Further, we found that the selection cassette had a considerable effect on the expression of the tagged protein; the remaining selection cassette reduced the expression level of the protein fused with fluorescent protein (Figs. 2 and 3). Moreover, recently, we have seen a reduction of the protein expression level after Cre-mediated selection cassette removal (38). This could be due to the effect of the 3’ UTR of the gene that was tagged by KI. For these reasons, removal of the selection cassette by Cre recombination is essential for the quantitative assessment of protein concentration. We quantified the endogenous protein concentrations of ERK2-mEGFP and RSK2-HaloTag-TMR by FCS (Fig. 6). The average concentration of ERK2-mEGFP measured in this study, 0.078 μM, was considerably lower than the ERK2 concentration reported previously, 0.68 μM for ERK2 in HeLa cells (14). This may have been due to the multiplication *of MAPK1* genes, because it is evident that the reduction of unlabeled EKR2 by KI of the *mEGFP* gene was quite limited (Fig. 2). The estimated total ERK2 concentration based on the mEGFP maturation efficiency and western blotting was 0.68 μM, which was consistent with the previous data. Therefore, the gene amplification, which is frequently observed in cancer cells, is a limitation of this technique. On the other hand, the RSK2-HaloTag-TMR concentration, 0.097 μM, was highly similar to the previously reported RSK2 concentration of 0.15 μM (14) (Fig. 3). The estimated total RSK2 concentration from western blotting—namely, 0.50 μM—was comparable. Indeed, KI of the *HaloTag* gene reduced the endogenous RSK2 protein level by half or to zero (Fig. 3C and 3D), implying monoallelic- and biallelic KI, respectively. Intra-cellular variability is a technical challenge in the quantification of protein concentration by FCS. The heterogeneity of the subcellular environment, including heterogeneity among the cellular organelles, causes the intra-cellular variability, since a positive correlation was observed between the protein abundance of ERK2-mEGFR and RSK2-HaloTag-TMR (Fig. 6B). FCS measures fluorescence fluctuation in a tiny confocal volume (~1 fL), and therefore the intra-cellular variability is unavoidable. For accurate determination of the protein concentrations in the cells, the intracellular variability can be canceled out by averaging repeated measurements at different points in each cell. Nonetheless, photobleaching still occurs in this approach and must be minimized.

We also quantified *in vivo K_d_* values for the binding of ERK2-mEGFP and RSK2-HaloTag-TMR by FCCS (Fig. 6). The average *in vivo K_d_* value measured in endogenously-tagged ERK2 and RSK2 in this study was 0.63 μM (Fig. 6C), but this value was higher than the *in vitro K_d_* value of ERK2 and RSK1 binding (39). This difference could be due to competitive bindings from endogenous proteins (14). Interestingly, the *in vivo K_d_* value measured in this study was lower than the *in vivo K_d_* value of 1.3 μM in overexpressed ERK2-mEGFP and RSK2-HaloTag (14). PEA-15 is known to function as a scaffold to enhance RSK2 activation by ERK (40). Thus, in the previous study, the overexpressed amounts of ERK2-mEGFP and RSK2-HaloTag might have exceeded the amount of endogenous PEA-15, resulting in an overestimation of the *in vivo K_d_* values. Further, the *in vivo K_d_* values of ERK2-mEGFP and RSK2-HaloTag-TMR binding were hardly correlated with the expression levels of those proteins (Fig. 6D), suggesting that the binding between ERK2 and RSK2 fluctuates in a manner dependent on unknown mechanisms, such as competitive binding, molecular crowding, posttranslational modification and/or scaffold proteins. In the future, quantitative analysis of the hetero-trimer complex could be a fascinating way to verify the scaffolding function of PEA-15 for ERK2-RSK2 binding (41).

Following EGF stimulation, ERK2-mEGFP was transiently dissociated from RSK2-HaloTag-TMR in the cytosol (Fig. 6G). It has been reported that ERK binding is differently regulated among RSK isoforms; RSK1 dissociated completely from ERK1/2 upon EGF stimulation, RSK2 only partially dissociated from ERK1/2, and RSK3 remained bound to ERK1/2 (27). These previous findings are consistent with our finding that a part of the complex of ERK2- mEGFP and RSK2-HaloTag-TMR was dissociated 10 min after EGF stimulation, while the other part remained (Fig. 6). The finding that the activation of RSK2 upon EGF stimulation is comparable to that of RSK1 suggested that the phosphorylation of RSK2 by ERK1/2, PDK1 and/or autophosphorylation (27, 42) reduced the affinity to ERK2 to some extent.

In summary, we established a method for quantifying the endogenous protein concentration and the *K_d_* value in a living cell. Further studies will be needed to improve the KI efficiency, the brightness of fluorescent proteins, and the quantification of proteins localized at specific cellular compartments such as the plasma membrane.

## Experimental procedures

### Design of gRNAs of CRISPR/Cas9 for gene KI

pX459 (pSpCas9(BB)-2A-Puro) was a gift from Feng Zhang (Addgene plasmid #62988). All guides are designed to overlap the stop codon of *MAPK1* or *RSK2*, so that these guide RNAs do not recognize the target sequences after KI. Guide sequences (without PAM sequence) were as follows: for *MAPK1*, TCTTAAATTTGTCAGGTACC; for *RSK2*, CTCACTGAGGTCACTTCACA.

### Construction of KI donor vectors

Truncated-thymidine kinase (dTK) originated from pPB-CAG.OSKM-puDtk (a gift from Kosuke Yusa and Allan Bradley (43)) and the neomycin-resistance gene (neo) were conjugated by PCR (Fig. S1). KOD FX neo (TOYOBO, Osaka, Japan) was used for all PCR reactions. Homology arms of the HDR vector were amplified from genomic DNA of HeLa cells extracted with QuickExtract DNA Extraction Solution (Epicentre, Madison, WI) and attached to an insertion cassette by PCR. 40 bp homology arms for the MMEJ donor were attached by PCR.

### Cell culture

HeLa cells were purchased from the Human Science Research Resources Bank (Sennanshi, Japan). HEK-293T cells were obtained from Invitrogen as Lenti-X 293 cells (Invitrogen, Carlsbad, CA). HeLa cells and HEK-293T cells were maintained in DMEM (Wako, Osaka, Japan) supplemented with 10% FBS. For imaging, HeLa cells were plated on 35-mm glass-base dishes (Asahi Techno Glass, Tokyo). At least 3 h before observation, HeLa cells were maintained with FluoroBrite DMEM (LifeTechnologies, Carlsbad, CA) supplemented with 1% GlutaMAX (LifeTechnologies).

For HaloTag imaging, the cells were incubated with 100 nM HaloTag-tetramethylrhodamine (TMR) for at least 18 h. After that, the cells were washed twice with PBS, then incubated with FluoroBrite DMEM supplemented with 1% GlutaMAX for 30 min. The medium was again replaced with the same medium.

### Establishment of the KI cell line

HeLa cells were plated on a 24-well plate, and transfected with 1 μg pX459 vectors and 50 ng KI donor DNA by using Polyethyleneimine “Max” MW 40,000 (Polyscience Inc., Warrington, PA), followed by puromycin selection. Three days after transfection, transfected cells were seeded onto 35-mm dishes and treated with 0.5 mg/ml G-418 (Invivogen, San Diego, CA) for more than 10 days. The genomic DNAs of these cells were extracted with QuickExtract DNA Extraction Solution (Epicentre), and analyzed by PCR.

To isolate MAPK1-EGFP KI cells, the selected cells were analyzed and sorted by FACSaria IIu (BD Biosciences, San Jose, CA). mEGFP fluorescence was detected using 488 nm laser light and a 518-548 nm BP emission filter, followed by sorting of mEGFP-positive cells and limiting dilution for single cell cloning. For single cell isolation of RSK2-HaloTag KI cells, G418-selected cells were subjected to the limiting dilution method.

### Adeno-associated viruses expressing Cre and removal of selection marker

The cDNA of Cre was inserted into pAAV-MCS (Cell Biolabs Inc., San Diego, CA), generating pAAV-Cre. HEK-293T cells were co-transfected with pAAV-Cre, pAAV-DJ, and pHelper to produce recombinant AAV expressing Cre recombinase. Three days after transfection, HEK-293T cells were collected and resuspended in 1 mL DMEM, followed by 4 freeze-thaw cycles. After the final thaw and centrifugation, 10 μL supernatant was added to each well of a 24-well plate to remove the selection marker. At least 6 days after AAV-Cre infection, the cells were selected with 50 μM Ganciclovir (Wako).

### Immunoblotting

Cells were lysed in 2x SDS sample buffer. After sonication, the samples were separated by 5-20% or 7.5% SDS-polyacrylamide gel electrophoresis (Nacali Tesque Inc., Kyoto, Japan), and transferred to polyvinylidene difluoride membranes (Millipore, Billerica, MA). After blocking with Odyssey Blocking Buffer (TBS) (LI-COR Biosciences Inc., Lincoln, NE) for 1 h, the membranes were incubated with primary antibodies diluted in blocking buffer, followed by secondary antibodies diluted in blocking buffer. Fluorescence levels were detected by an Odyssey infrared scanner (LI-COR Biosciences Inc.).

The following antibodies were used in this study: anti-green fluorescent protein (GFP) antibody (632375; Clontech, Palo Alto, CA), anti-p44/p42 MAP kinase (ERK1/2) antibody (4695; Cell Signaling Technology, Danvers, MA), anti-RSK2 (sc-9986; Santa Cruz Biotechnology, Santa Cruz, CA), anti-HaloTag antibody (G9281; Promega, Madison, WI), anti-α-tubulin (PM054, MBL), IRDye680LT goat anti-rabbit IgG antibody (no. 925-68021; LI-COR), and IRDye800CW donkey anti-mouse IgG antibody (no. 925-32212; LI-COR).

### Imaging by confocal microscope

Cells were imaged with a laser scanning confocal microscope (FV1200; Olympus) equipped with gallium arsenide phosphide (GaAsP) detectors. The excitation lines were set at 473 nm and 559 nm. The excitation beam was reflected by a DM 405/488/559 dichroic mirror and focused by an oil immersion objective lens (Uplsapo 60XO, NA 1.35; Olympus). The emitted light was detected through a bandpath filter with wavelengths of 495 to 540 nm for mEGFP and 575 to 630 nm for HaloTag-TMR.

### FCS and FCCS

FCS and FCCS data were obtained as described previously (14). Briefly, cells were imaged with a laser scanning confocal microscope (FV1000 or FV1200; Olympus) equipped with GaAsP detectors. The excitation lines and power were set at 473 nm (0.1%) and 559 nm (0.1%). The excitation beam was reflected by a DM 405/488/559 dichroic mirror and focused by an oil immersion objective lens (Uplsapo 60XO, NA 1.35; Olympus). Time-series data of fluorescence fluctuation were obtained by point scan (2.0 μsec/1 sampling) in photon count mode, and total time-series data were typically 8,000,000 points. Of note, first 1,600,000 data points were used for the FCS and FCCS analysis to exclude the effect of photobleaching.

FCS and FCCS data were analyzed as follows. First, in order to correct the intensity decay due to photobleaching, we scaled intensity traces so that the corrected traces have constant means and fluctuations in time (See Eqs. 21 and 22 in Ries et al. (44)). Then, we used the multiple tau Python package (Version 0.1.9) developed by Paul Müller (https://pypi.org/project/multipletau/, Accessed 9 Jan, 2018) to compute auto- and cross-correlations. The numerical correlation functions were fitted by the theoretical model:

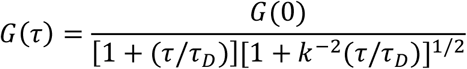

where τ, *τ_D_, k*, and *G*(0) are the time delay, the correlation time, the structure parameter, and the amplitude of correlation function, respectively. Note that *G*(0) is related to fluorophore concentration *C* as

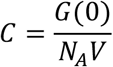

where *N*_A_ is the Avogadro number and *V* is the effective confocal volume. The values of *k* and *V* were determined in each microscopy setup by the common procedure using Rhodamine 6G solution (14), and typical values were 5-5.8 and 0.3-0.59 fL, respectively. We used the “curve_fit” function in Python’s SciPy package for fitting, and thereby obtained the concentrations of mEGFP, Halotag-TMR, and the complex. In addition, we corrected the concentration of the complex as described in a previous study (14).

To estimate the dissociation constant *K_d_*, we adopted the following equations:

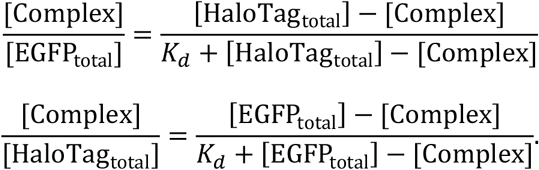

The *in vivo K_d_* value was obtained by averaging the values derived from these equations.

### Estimation of total ERK2 and RSK2 concentration

Total concentrations of ERK2 and RSK2 in a HeLa cell were estimated as follows. First, the average concentrations of ERK2-mEGFP and RSK2-HaloTag-TMR were 0.078 μM and 0.097 μM in HeLa/ERK2-mEGFP/RSK2-HaloTag (MMEJ#10/PITCh#3) as measured by FCS (Fig. 6B). Next, we obtained an mEGFP fluorophore maturation efficiency of 0.92 (Fig. 5C). The HaloTag-TMR labeling efficiency was assumed to be 100%. Finally, western blotting data (Fig. 2D for ERK2 and Fig. 3D for RSK2) provided the ratio value of ERK2 to ERK2-mEGFP, which was 8.0 and that of RSK2 to RSK2-HaloTag, which was 5.1. ERK1 and tubulin were used as loading controls for ERK2 and RSK2, respectively. Based on these values, the total ERK2 concentration was estimated as follows: 0.078/0.92*8.0 = 0.68 μM. The total RSK2 concentration was estimated as follows: 0.097*5.14 = 0.50 μM.

## Acknowledgement

We thank the members of the Matsuda Laboratory and Aoki Laboratory for their helpful input. This work was supported by the Spectrography and Bioimaging Facility and Functional Genomics Facility, NIBB Core Research Facilities. A. Kawagishi, K. Hirano, N. Nishimoto, E. Ebine, and K. Onoda are also to be thanked for their technical assistance.

## Author contributions

A.T.K., Y.G., and K.A. designed and performed the experiments. A.T.K., Y.G., Y.K. analyzed the data. A.T.K., Y.G., Y.K, M.M., and K.A. wrote the manuscript. A.T.K. and Y.G. contributed equally to this work.

## FOOTNOTES

K.A. and M.M. were supported by the Platform for Dynamic Approaches to Living System from the Ministry of Education, Culture, Sports, and Science, Japan, and by JST, CREST Grant No. JPMJCR1654. K.A. was supported by JSPS KAKENHI Grants No. 16H01425 “Resonance Bio”, 16H01447, 16KT0069, 18H04754 “Resonance Bio”, and 18H02444, and by funds from the Hori Sciences and Arts Foundation and the Nakajima Foundation. M.M. was supported by MEXT/JSPS KAKENHI Grant No. 15H05949 “Resonance Bio”. A.T.K. was supported by a JSPS Grant-in-Aid for Research Fellows.

## Supporting Information

**Figure S1.**
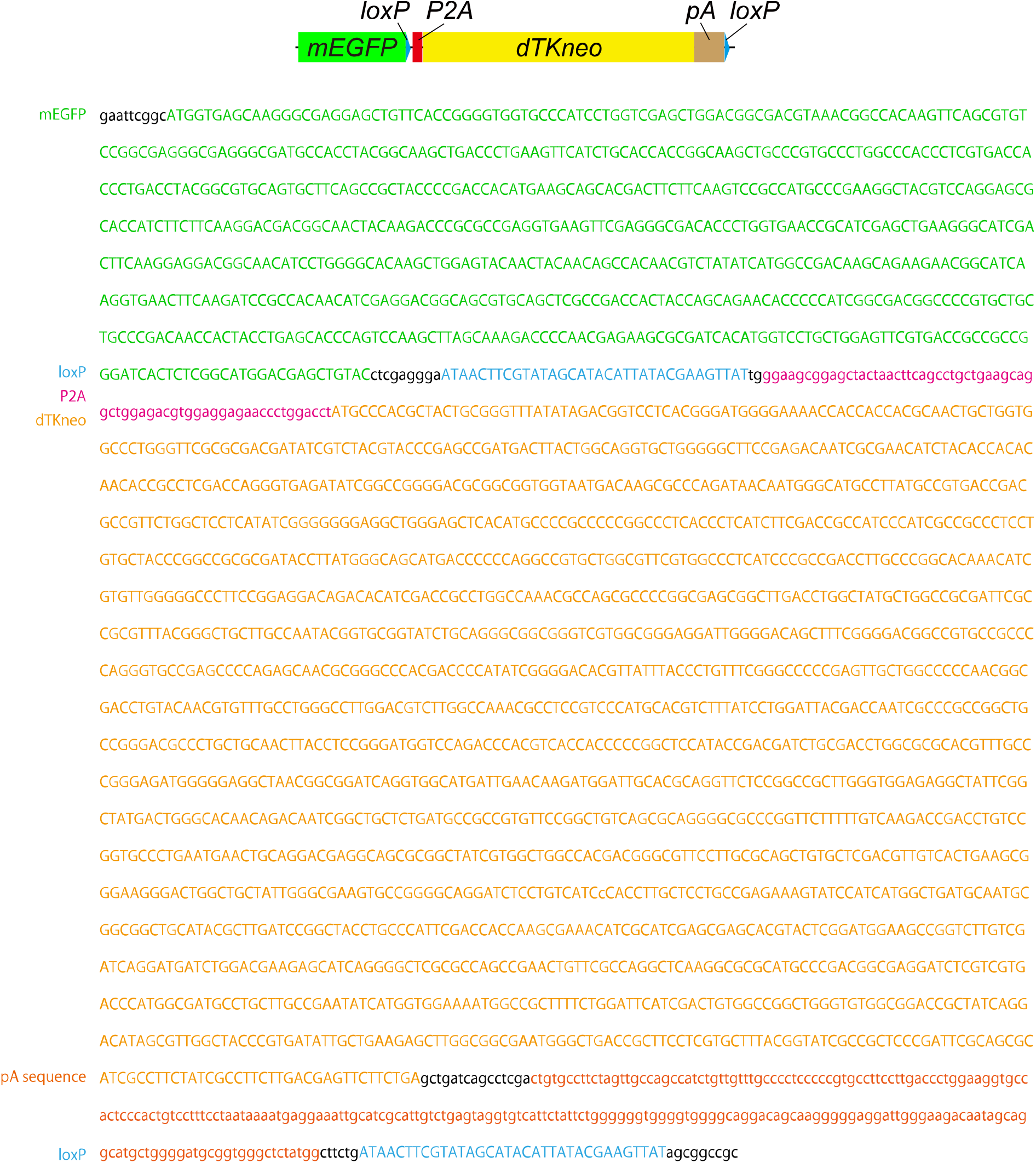
DNA sequence of the selection cassette for the KI vector. Schematic representation (upper) and DNA sequence (lower) of the selection cassette. The vector sequence and map are available in https://benchling.com/s/seq-Fk3xGD0OazVzjB1UHQBI.

**Figure S2.**
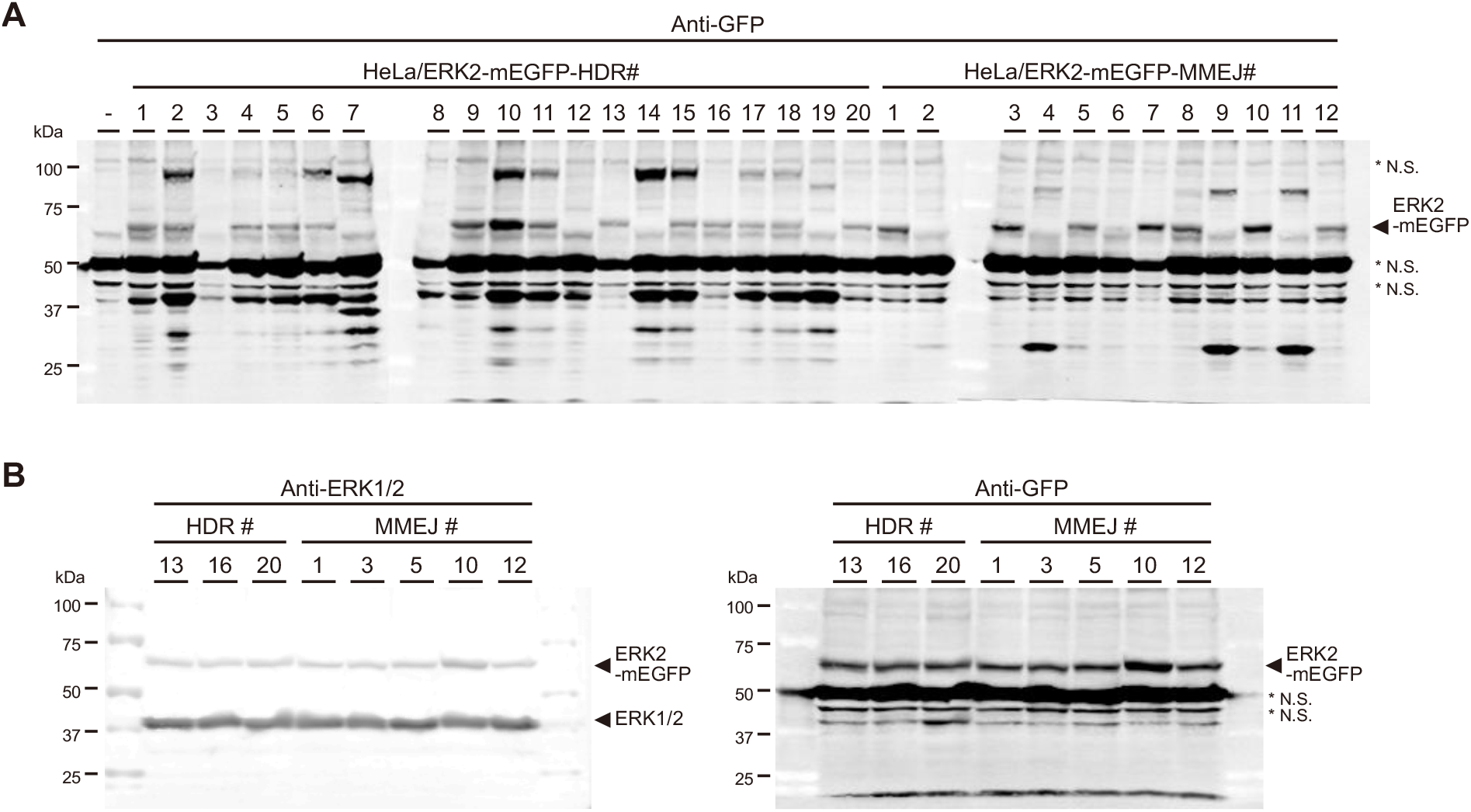
Validation of mEGFP knock-in at the human hMAPK1 locus. (A) The cell lysates were obtained from parental HeLa cells (-), and the indicated individual clones from HeLa/ERK2-mEGFP-HDR or HeLa/ERK2-mEGFP-MMEJ in Figure 2C, and subjected to immunoblotting with anti-GFP antibody. The arrowhead indicates ERK2-mEGFP. An asterisk (*) indicates a nonspecific (N.S.) signal. (B) The cell lysates from the indicated clones were analyzed by immunoblotting with anti-ERK1/2 (left) and anti-GFP (right) antibodies.

**Figure S3.**
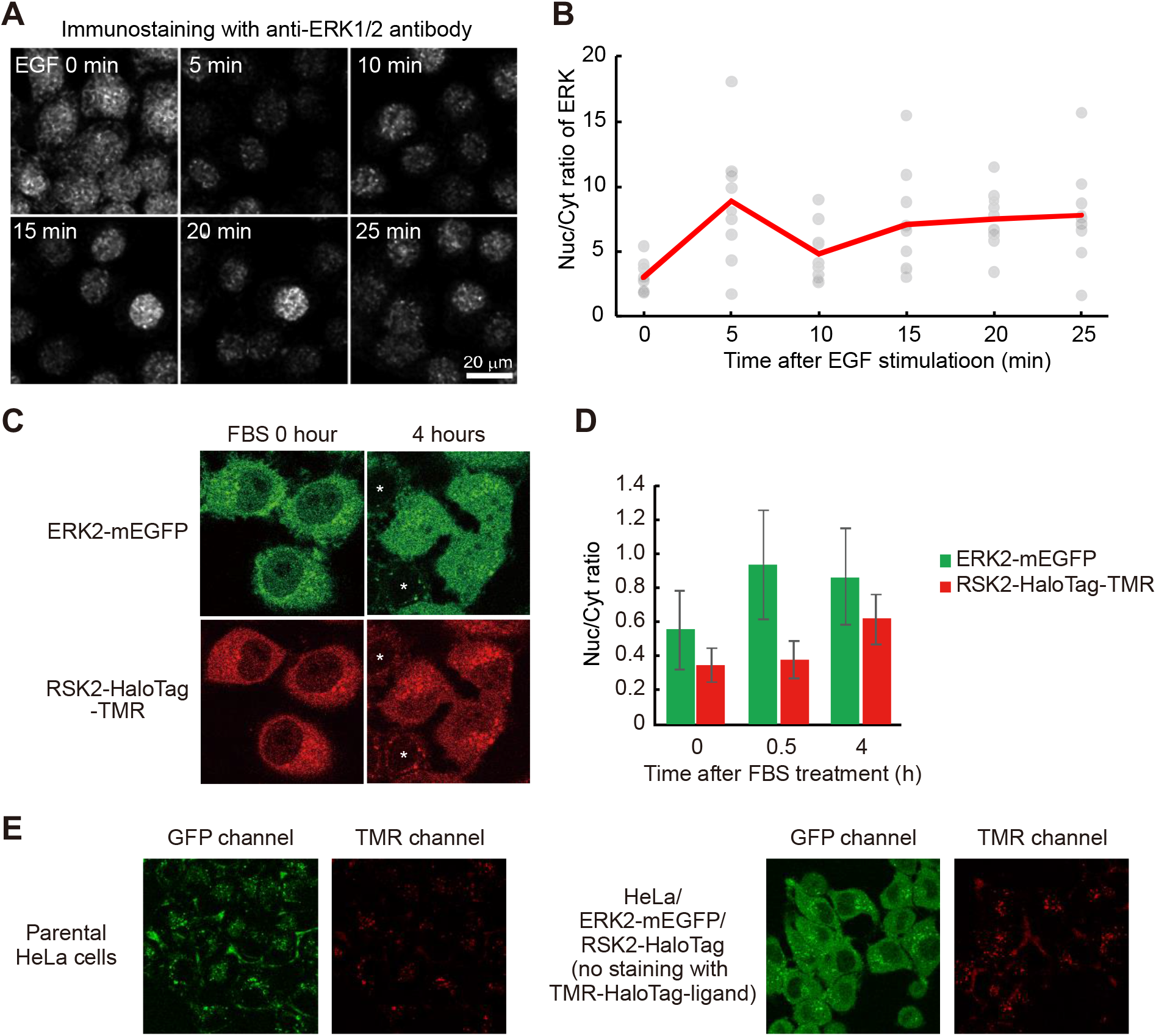
Nuclear accumulation of endogenous ERK2-mEGFP and RSK2-HaloTag-TMR upon stimulation. (A and B) HeLa cells were stimulated with 10 ng/mL EGF for the indicated amount of time, followed by fixation. The cells were subjected to immunofluorescence staining with anti-ERK1/2 antibodies. The representative images are shown (A). The ratio of fluorescence intensity in the nucleus to that in the cytoplasm was quantified and plotted as a function of time after EGF stimulation (B). The red line indicates the averaged data. (C and D) HeLa/ERK2-mEGFP-MMEJ#10/HaloTag-PITCh#3 cells and parental HeLa cells were cocultured. The cells were stimulated with 10% FBS and imaged after 4 h. The representative images are shown (C). An asterisk (*) indicates a parental HeLa cell. The averaged ratio value of fluorescence intensity in the nucleus to that in the cytoplasm was quantified and plotted as a bar graph with the SD (D). N = 10 cells. (E) The representative auto-fluorescences in the GFP and TMR channel in HeLa cells are shown.

**Figure S4.**
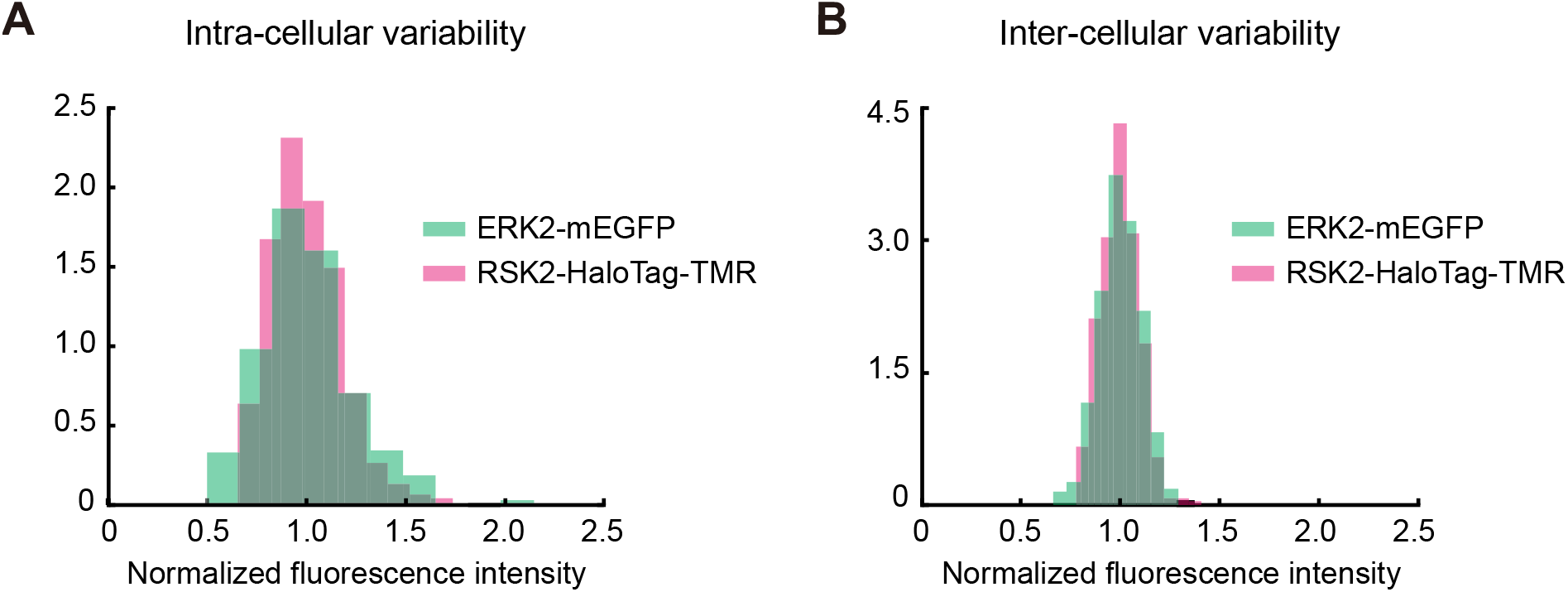
Comparison of intra- and intercellular variability of ERK2-mEGFP and RSK2-HaloTag-TMR. ERK2-mEGFP and RSK2-HaloTag-TMR images of HeLa/ERK2-mEGFP-MMEJ#10/HaloTag-PITCh#3 cells were obtained by a confocal microscope. (A) The fluorescence intensities of 10 points in the cytoplasmic area were arbitrarily picked up in a single cell and averaged to obtain the normalized fluorescence intensity. The normalized fluorescence intensity is plotted as a histogram. N = 42 cells. CV (ERK2-mEGFP) = 0.232, CV (RSK2-HaloTag-TMR) = 0.179. (B) The fluorescence intensity in a whole cell area was measured and averaged among cells to calculate the normalized fluorescence intensity. The normalized fluorescence intensity is plotted as a histogram. N = 42 cells. CV (ERK2-mEGFP) = 0.106, CV (RSK2-HaloTag-TMR) = 0.097.

**Figure S5.**
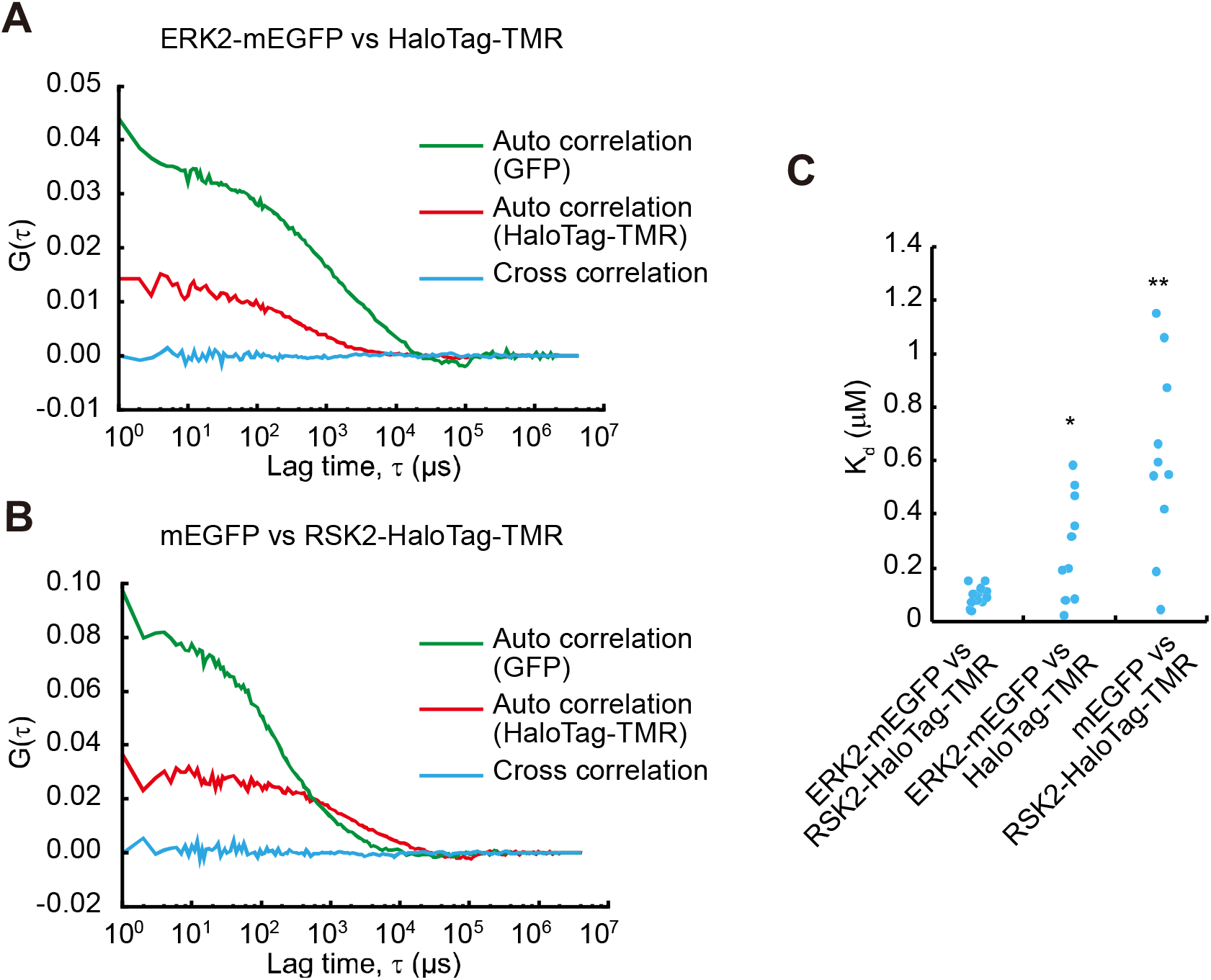
Negative control experiments of ERK2 and RSK2 binding. (A and B) HeLa cells exogenously expressing ERK2-mEGFP and HaloTag-TMR (A) or mEGFP and RSK2-HaloTag-TMR (B) were subjected to FCCS analysis. The representative correlation curves are shown. (C) The K_d_ values are plotted in the indicated binding pairs. N = 17, 10, 10 cells. \**p* < 0.05, \*\**p* < 0.01 by Welch’s *t*-test.

